# Rostral ventromedial medulla (RVM) projects to the lateral hypothalamic area (LHA) to drive aversion and anxiety

**DOI:** 10.1101/2024.11.29.626042

**Authors:** Mousmi Rani, Devanshi Piyush Shah, Arnab Barik

**Affiliations:** Center for Neuroscience, Indian Institute of Science, Bengaluru, India, 560012

## Abstract

Neurons in the LHA are critical drivers of behavioral and physiological responses to acute and chronic stress. However, the roles of the specific pre-synaptic inputs to the LHA in driving stress and resultant physiological effects are yet to be fully understood. Here, taking advantage of mouse viral genetics, rabies tracing, optogenetics, chemogenetics, and fiber photometry, we show that the excitatory projections from the RVM to LHA drive stress-induced anxiety. This is a surprising finding since, traditionally, RVM has been studied in the context of opioidergic pain modulation through its inhibitory projections to the spinal cord. We find that the LHA neurons receiving inputs from the RVM, when activated, do not alter the nociceptive thresholds yet are sufficient to drive anxiety-like behaviors. These LHA neurons are recruited by acute restraint, which is known to cause stress. On the other hand, the LHA-projecting RVM neurons are responsive to both noxious thermal stimuli and acute restraint, promoting stress-induced anxiety, yet with no effect on pain thresholds. Together, we found an ascending neural pathway between RVM and LHA that mediates stress-induced anxiety.

**Significance statement:** There is a strong correlation between pain and anxiety. However, the underlying neural mechanisms are poorly understood. Here, we reveal a novel neural pathway between the rostral ventromedial medulla (RVM) and lateral hypothalamus (LHA) that can potentially convert painful experiences into stress and anxiety. The traditional role of RVM is opioidergic modulation of pain. However, here we show that in addition to nociception, RVM neurons can play an essential role in the affective-motivational components of pain.

## Introduction

Stress and pain are inextricably linked (Lumley et al., 2011; Hannibal and Bishop, 2014). Acute stress increases pain thresholds, whereas chronic stress causes hyperalgesia (Terman et al., 1984; Amit and Galina, 1986; Butler and Finn, 2009). On the other hand, patients with chronic pain often suffer from stress and anxiety (Woo, 2010; De La Rosa et al., 2024). Thus, to develop better clinical interventions for chronic pain and comorbid anxiety, it is important to understand the neural mechanisms of how stress precipitates somatosensory hypersensitivity due to chronic pain and vice versa.

The excitatory neurons in the lateral hypothalamic area (LHA) mediate the expression of physiological and behavioral effects of stress and pain. Inputs from the lateral septum (LS) have been shown to inhibit a subpopulation of the excitatory LHA neurons upon restraint stress, and this inhibitory mechanism is essential for restraint-stress (RS) induced analgesia(Shah et al., 2024). Meanwhile, an independent excitatory neural population expressing parvalbumin (PV) and downstream projections to the periaqueductal gray (PAG) were found to be anti-nociceptive (Siemian et al., 2021). The VGlut2-expressing LHA neurons project to the lateral habenula and control a sex-specific aversive stress-like state in female mice (Calvigioni et al., 2023). The glutamatergic LHA neurons projecting to the lateral habenula responded to foot shocks and mildly aversive events (Lazaridis et al., 2019). In addition to the excitatory cells, the GABAergic Galanin-expressing neurons are activated by anxiogenic stimuli and are sufficient to lessen stress-induced anxiety (Owens-French et al., 2022). However, it remains unclear how the LHA neurons integrate noxious somatosensory and anxiogenic signals.

The stress-induced analgesia (SIA) and LHA-mediated pain modulation are dependent and occur through the downstream nociceptive neurons in the RVM (Willer et al., 1981; Mogil et al., 1996; Shah et al., 2024). Recent findings indicate that the RVM can be involved in chronic pain-associated depressive behaviors (Pagliusi et al., 2024). However, it is less clear how RVM may modulate the affective-motivational components of pain, such as anxiety. Here, we found that a subpopulation of nociceptive neurons in the RVM (RVM_pre-LHA_) projects to the LHA (LHA_post-RVM_). We found that the RVM_pre-LHA_ neurons and LHA_post-RVM_ are aversive and sufficient to drive mice to be anxious. Surprisingly, activation or inhibition of the RVM_pre-LHA_ or the LHA_post-RVM_ did not affect the nociceptive thresholds. Thus, our data indicate that in addition to modulating the nociceptive thresholds, the RVM neurons, through their LHA projections, can mediate the affective-motivational components of pain.

## Methods

### Animals

Animal care and experimental procedures were performed following protocols approved by the CPCSEA at the Indian Institute of Science, Bangalore. CD1 mice were housed at the IISc Central Animal Facility under standard animal housing conditions: 12 h light/dark cycle from 7:00 am to 7:00 pm with ad-libitum access to food and water; mice were housed in IVC cages in Specific pathogen-free (SPF) clean air rooms. All mice used in the behavioral assays were between 8 and 12 weeks old. All the behaviors were done during the light cycle.

### Viral vectors and stereotaxic injections

Mice were anesthetized with 2% isoflurane/oxygen before and during the surgery. An incision was made to expose the skull, and subsequently, the skull was aligned in a plane. Craniotomy was performed at the marked point using a hand-held micro-drill (RWD). A Hamilton syringe (10 ul) with a glass pulled needle infused 300 nL of viral particles (1:1 in saline) at 100 nL/min. The following coordinates were used to introduce the virus: LHA, AP: −1.70, ML: ±1.00, DV: −5.15; RVM, AP: −5.80, ML: +0.08, DV: −5.50; LS, AP: +0.50, ML: ± 0.25, DV: −2.50. Vector used and sources: pENN-AAV-hSynCre-WPRE-hGH (Addgene, Catalog# v181154-AAV1), pENN-AAV-hSynCre-WPRE-hGH (Addgene, Catalog# v173770-AAVrg), pAAV5-hsyn-DIO-eGFP (Addgene, Catalog# 50457-AAV1), pAAV5-FLEX-tdTomato (Addgene, Catalog# 28306-PHP.S), pAAV5-hsyn-DIO-hM3D(Gq)-mCherry (Addgene, Catalog# v141469), pAAV5-hsyn-DIO-hM4D(Gi)-mCherry (Addgene, Catalog# 44362-AAV5), AAV9-syn.flex.GcaMP6s (Addgene, Catalog# pNM V3872TI-R(7.5)), v651-1), AAV5-EF1a-DIO-hChR2-mCherry (Addgene, Catalog# 20297 Lot: v138748) AAV9-DIO-PSD95-TagRFP (Donated by Mark Hoon, NIH) AAV5-hSyn-DIO-mSyp1-eGFP(University of Zurich, Catalog# v484-9).

Tissue was harvested 21 days after viral injection for histology. Post-hoc histological examination of each injected mouse was used to confirm that viral-mediated expression was restricted to target nuclei.

### Optogenetic and Photometry Fiber Implantation

Fiber optic cannula from RWD; Ø1.25 mm Ceramic Ferrule, 200 μm Core, 0.22 NA, L = 6 mm were implanted at AP: −5.80, ML: -0.08, DV: −5.50 in the RVM and AP: −1.70, ML: ±1.00, DV: −5.15 in the LHA after AAV carrying GCamp6s or ChR2, were infused. Animals were allowed to recover for at least 3 weeks before performing behavioral tests. Successful labeling and fiber implantation were confirmed post hoc by staining for GFP/mCherry for viral expression and injury caused by the fiber for implantation. Animals with viral-mediated gene expression at the intended locations and fiber implantations, as observed in post hoc tests, were only included.

### Fiber Photometry

A dual-channel fiber photometry system from RWD (R820 model) was used to collect the data. The light from two light LEDs (410 and 470 nm) was passed through a fiber optic cable coupled to the cannula implanted in the mouse. Fluorescence emission was acquired through the same fiber optic cable onto a CMOS camera through a dichroic filter. The photometry data was analyzed using the RWD software, and .csv files were generated. The start and end of stimuli were timestamped. All trace graphs were plotted from .csv files using the GraphPad Prism software version 8.

### Behavioral Assay

A single experimenter handled behavioral assays for the same cohorts. Mice were acclimatized for at least 30 minutes in the behavior room before experiments. An equal male-to-female ratio was maintained in every experimental cohort and condition unless otherwise stated, with no significant differences seen between sexes in the responses recorded from the behavioral experiments. Wherever possible, efforts were made to keep the usage of animals to a minimum. Deschloroclozapine (DCZ) was diluted in saline (0.1 mg/kg) and injected intraperitoneal (i.p.) 15-20 minutes before behavioral experiments or histochemical analysis.

The thermal hotplate experiments were performed using Orchid Scientific Hot and Cold Plate Analgesiameter (HC-01). The specifications of the instrument used are- enclosure size: (L × W × H): 205 × 205 × 250 mm; plate size: (L × W × H): 190 × 190 × 06 mm; temperature range: −5°C to 60°C. On the experimental day, mice were placed on the 52°C plate temperature and recorded for one minute using three Logitech web cameras, placed at the left, right, and in front. Later, videos were quantified individually for mice’s nocifensive behavior (licks and shakes were scored). The tail flick analgesiometer (Orchid Scientific, India) was used according to the manufacturer’s instructions. The mice were restrained in a modified falcon tube, exposing only the tail. The mouse’s tail was then subjected to heat via an intense light beam, and the time the mouse flicked its tail was recorded.

A fiber-coupled laser (channelrhodopsin activation; RWD, China) was used for optogenetic stimulations. Before the behavioral testing, the optic fibers were connected to the cannula implanted in the brain of the mice. The animals were habituated for 30 minutes before the commencement of the experiments. The light-dark box tracking and estimations of the time spent in either chamber were carried out with DeepLabCut, and the data were trained and analyzed on a custom-built computer system with AMD Ryzen 9 5900x 12-core processor 24 with NVIDIA Corporation Graphics Processing Unit (GPU) (Mathis et al., 2018).

For GCaMP fiber photometry recordings, the mice were subjected to five primary behavioral assays. In the immobilization experiments, the experimenter physically restrained the mice by pressing them down by hand for approximately 10 seconds. In the tail suspension experiments, the mice were suspended upside down by their tail for 10 seconds. In the tail-flick assay, photometry signals were recorded through the fiber-coupled cannulae that passed through a modified falcon tube to allow unrestricted recording. In the tail-pinch experiment, the animal’s tail was pinched with forceps. On the hot plate test, the mice were acclimatized to the equipment with the optic fiber connected to the cannulae a day before experimentation. The equipment was first allowed to reach the desired temperature during the experiments, and then the animals were introduced to the hot plate test.

### Immunostaining, multiplex in situ hybridization, and confocal microscopy

Mice were anesthetized with isoflurane and perfused transcardially with 1X PBS (Takara) and 4% PFA (Ted Pella, Inc.) consecutively for immunostaining experiments. Tissue sections were rinsed in 1X PBS and incubated in the blocking buffer (5% Bovine Serum Albumin; 0.5% Triton X-100; PBS) for 1 hour at room temperature. Sections were incubated in the primary antibody in the blocking buffer at room temperature overnight. Sections were rinsed 1-2 times with PBS and incubated for 2 hours in Alexa Fluor conjugated goat anti-rabbit/ chicken or donkey anti-goat/rabbit secondary antibodies (Invitrogen) at room temperature, washed in PBS, and mounted in VectaMount permanent mounting media (Vector Laboratories Inc.) onto charged glass slides (Globe Scientific Inc.). Subsequently, sections were imaged on the upright fluorescence microscope (Khush, Bengaluru) (2X, 4X, and 10X lens) and ImageJ/FIJI image processing software to process the images.

Fresh brains were harvested for in situ hybridization experiments. Multiplex ISH was done with a manual RNAscope assay (Advanced Cell Diagnostics). Probes were ordered from the ACD online catalog. For the anatomical studies, the images were collected with 10X and 20X objectives on a laser scanning confocal system (Leica SP8 Falcon) and processed using the Leica image analysis suite.

### Quantification and Statistical Analysis

All statistical analyses were performed using GraphPad PRISM 8.0.2 software. ns > 0.05, ∗ p ≤ 0.05, ∗∗ p ≤ 0.01, ∗∗∗ p ≤ 0.001, ∗∗∗∗ p ≤ 0.0005.

## Results

### LHA and RVM are bi-directionally connected

Excitatory neurons of the LHA, receiving inhibitory LS inputs, project to the RVM to facilitate stress-induced analgesia (Shah et al., 2024). However, the axon terminals of RVM neurons enriched with synaptic vesicles were found in the LHA (Shah et al., 2024). Implying that the RVM and LHA are bidirectionally connected. Thus, we wondered if the neurons in the LS-LHA-RVM pathway overlap with the RVM-LHA circuitry. To that end, we stereotaxically injected the anterograde transsynaptic AAV1-hSyn-FlpO or AAVTranssyn-FlpO(Zingg et al., 2017, 2022) in the LS and AAV1-hSyn-Cre or AAVTranssyn-Cre in the RVM (Figure 1A). We intersected the FlpO and Cre recombinases in the LHA with the FlpO-dependent fDIO-tdTomato and DIO-GFP. This strategy allowed us to simultaneously label the neurons post-synaptic to LS with tdTomato and post-synaptic to RVM with GFP (Figure 1B-D). We found that small numbers of tdTomato and GFP-expressing LHA neurons overlapped (Figure 1E). Hence, the RVM-LHA circuitry may function independently of the LS-LHA-RVM pathway for stress-induced analgesia elucidated before (Shah et al., 2024).

**Figure 1:**
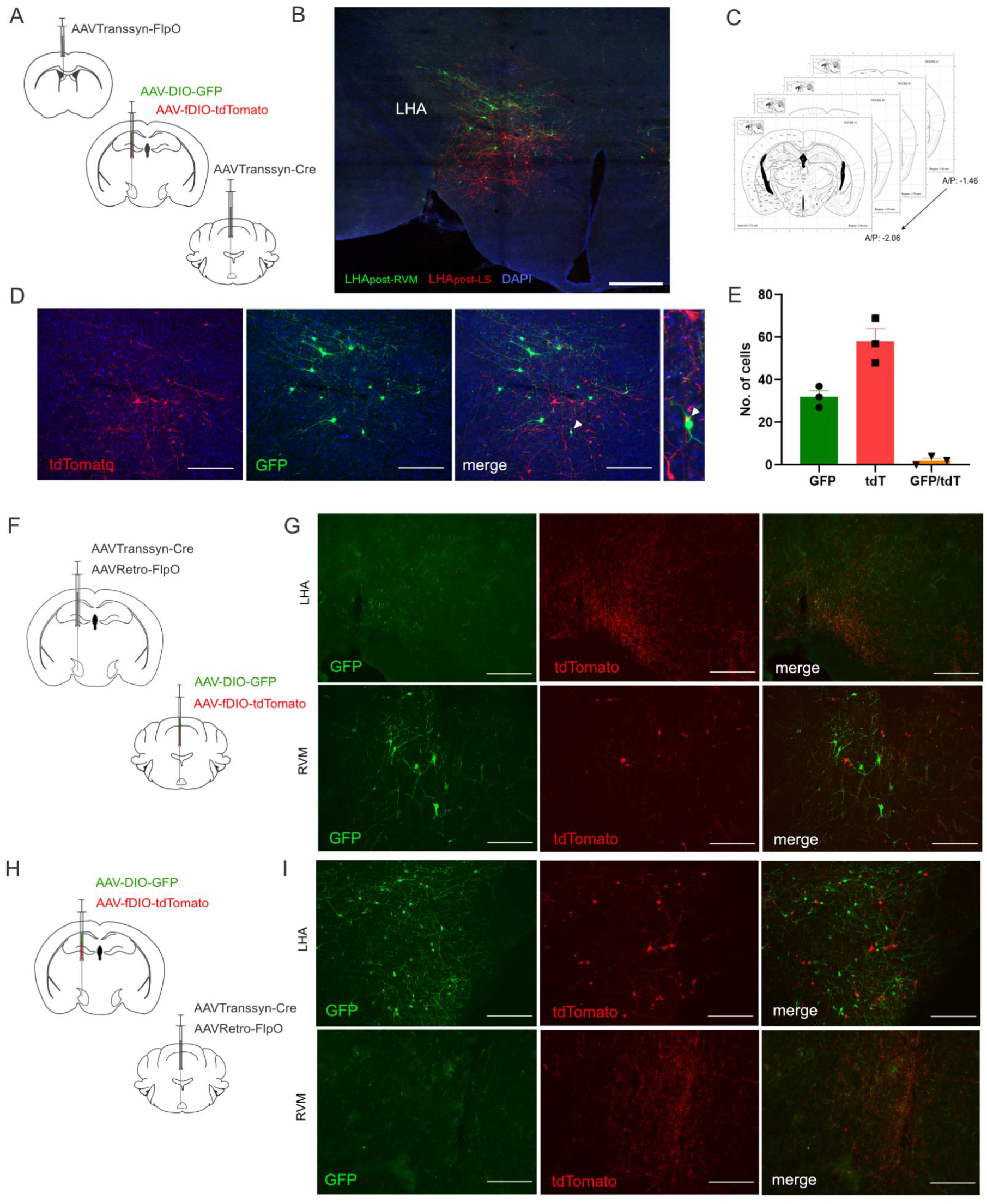
RVM and LHA are bidirectionally connected. (A) AAVTranssyn-FlpO in the LS; AAVTranssyn-Cre in the RVM; DIO-eGFP and fDIO-tdTomato in the LHA were stereotaxically injected. (B) Expression of Cre-dependent GFP and Flp-dependent tdTomato in the LHA. (C) The mouse brain figure has been reproduced from the Allen mouse brain atlas denoting the A/P axis range (AP:-1.46 to AP: −2.06) considered for cell counting. (D) Representative images show GFP-labeled LHA_post-RVM_ (green) and tdTomato-labeled LHA_post-LS_ (red) neurons. (E) Bar graph representing the quantification of LHA_post-RVM_ (green, 33.33 ± 3.18), LHA_post-LS_ (red, 57.33 ± 5.61), and co-localized (yellow, 3.00 ± 1.15) neurons in the LHA (n=3). (F) AAVTranssyn-Cre and AAVRetro-FlpO were injected in the LHA; DIO-GFP and Flp-tdTomato were injected in the RVM of wild-type mice. (G) Representative image of GFP (green) and tdTomato (red) expressions in the RVM. Non-overlapping expression of axon terminals in the LHA. (H) AAVTranssyn-Cre and AAVRetro-FlpO were injected in the RVM; DIO-eGFP and Flp-tdTomato were injected in the LHA of wild-type mice. (I) Representative image of GFP (green) and tdTomato (red) expressions in the LHA. Non-overlapping expression of axon terminals in the RVM. Green, GFP; Red, tdTomato; Blue, DAPI. Scale bars, 100 μm.

Next, we attempted to determine if the RVM neurons that receive inputs from the LHA are the same as the ones that innervate the LHA. We injected AAVTranssyn-Cre and retrogradely transported AAVRetro-FlpO (Tervo et al., 2016) in the LHA, DIO-GFP, and fDIO-TdTomato in the RVM (Figure 1F). This experiment allowed us to label the RVM neurons receiving inputs from LHA (RVM_post-LHA_) in GFP and RVM neurons projecting to the LHA (RVM_pre-LHA_) in tdTomato (Figure 1G). We found that the GFP and TdTomato are expressed in a non-overlapping group of cells in the RVM. Similarly, we injected the AAVTranssyn-Cre and AAVRetro-FlpO in the RVM, intersecting with DIO-GFP and fDIO-TdTomato in the LHA (Figure 1H). Here, we found that the neural population in the LHA that innervates RVM (LHA_pre-RVM_) and receives mono-synaptic inputs from the RVM (LHA_post-RVM_) are independent (Figure 1I). Together, these findings indicate that the RVM-LHA are bi-directionally connected. However, there are two distinct directions of the flow of information; in one, the LS-LHA-RVM circuitry facilitates SIA, whereas, in another, the RVM-LHA circuitry serves yet unknown functions.

### RVM neurons synapse onto Vglut2-expressing LHA neurons

LHA is mainly comprised of inhibitory (GABAergic) neurons, with a relatively minor population of excitatory (glutamatergic) cells. The glutamatergic neurons are involved in SIA and innervate the PAG and RVM to modulate nociceptive thresholds(Siemian et al., 2020, 2021; Shah et al., 2024). Here, we asked if the LHA_post-RVM_ neurons are glutamatergic. To that end, we injected the AAVTranssyn-Cre in the RVM and DIO-TdTomato in the LHA, allowing us to label the LHA_post-RVM_ neurons with the tdTomato fluorescent protein (Figure 2A). Next, we performed fluorescent multiplex in-situ hybridization (RNAscope) (Wang et al., 2012) to determine if the tdTomato expressing LHA_post-RVM_ neurons co-express *slc17a6* mRNA. Since *slc17a6* mRNA encodes the gene for glutamate transporter, VGlut2, and the LHA_post-RVM_ neurons express both *slc17a6* and *tdtomato*, we concluded that the LHA_post-RVM_ neurons are glutamatergic (Figure 2B). Next, we wondered if the RVM_pre-LHA_ neurons are glutamatergic or GABAergic. We labeled the RVM_pre-LHA_ neurons with tdTomato by delivering AAVRetro-Cre in the LHA and DIO-tdTomato in the RVM (Figure 2C). Simultaneous RNAscope for *slc17a6, slc32a1* (gene encoding the GABA-transporter, Vgat), and *tdtomato* showed that the RVM_pre-LHA_ neurons are glutamatergic (Figure 2D). Thus, the RVM_pre-LHA_ neurons and LHA_post-RVM_ are glutamatergic, forming an excitatory-excitatory circuitry. To further confirm this finding, we expressed GFP fused with mouse synaptophysin (mSyp1-eGFP) (Barik et al., 2021) in a Cre-dependent manner in the VGlut2-expressing neurons in the RVM (Figure 2E). This allowed the expression of GFP in the synaptic vesicles of the excitatory RVM neurons, which included the RVM_pre-LHA_ neurons.

**Figure 2:**
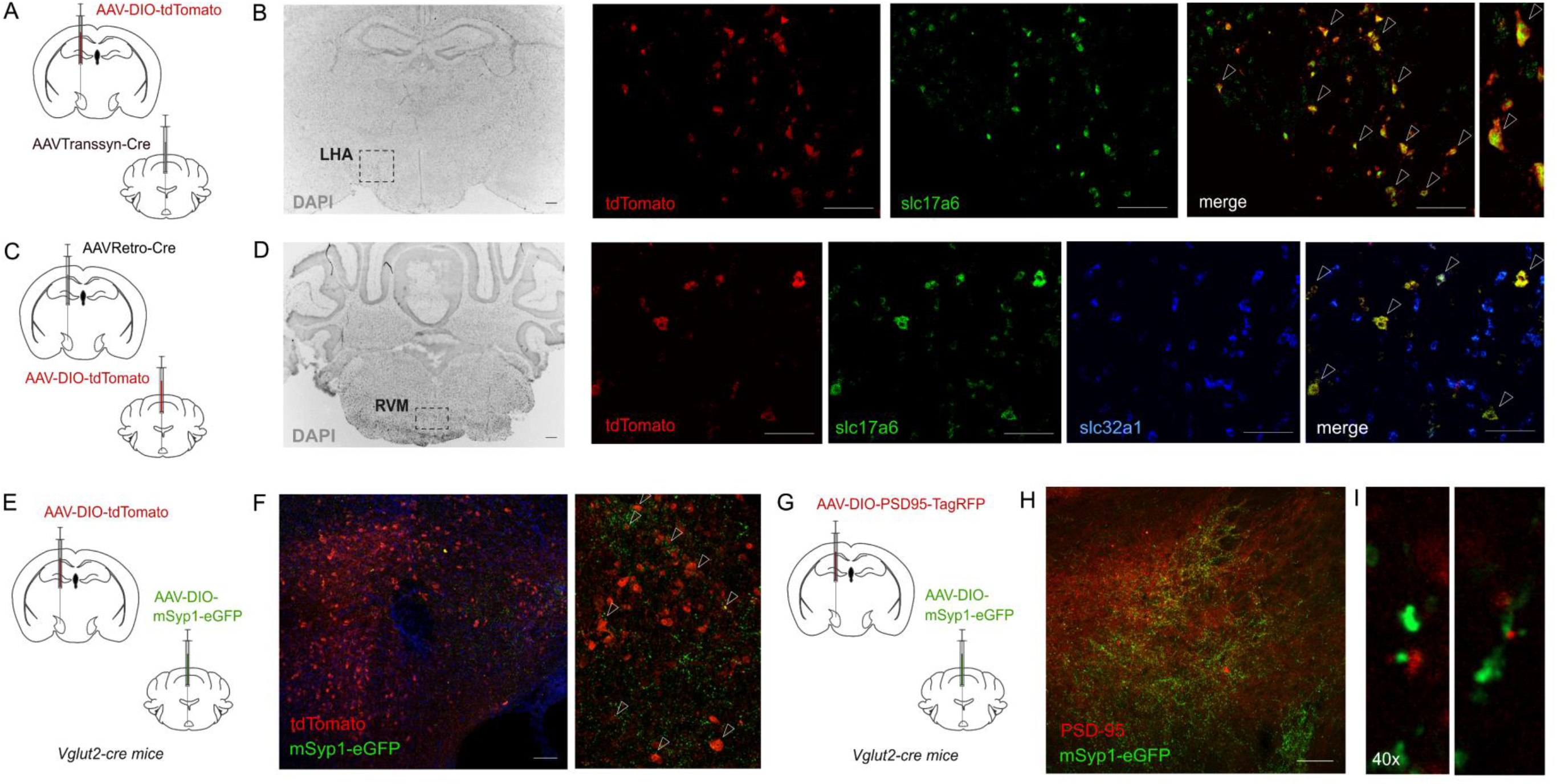
RVM forms excitatory synapses with the LHA neurons. (A) AAVTranssyn-Cre in the RVM and DIO-tdTomato in the LHA were injected in wild-type mice. (B) In-situ hybridization shows co-localization (yellow) of LHA_post-RVM_ neurons (red) with *slc17a6* expression (green). Red, tdTomato; Green, slc17a6; Grey, DAPI. Scale bars, 100 μm (C) AAVRetro-Cre in the LHA and DIO-tdTomato in the RVM were injected in wild-type mice. (D) Using in situ hybridization, the RVM was labeled with probes against *tdtomato, slc17a6,* and *slc32a1*. Yellow represents the co-localization of tdTomato-positive (red) and slc17ac (green) cells in the RVM. Red, *tdtomato*; Green, slc17a6; Blue, *slc32a1*; Grey, DAPI. Scale bars, 100 μm. (E) DIO-tdTomato in the LHA and DIO-mSyp1-eGFP in the RVM were injected in VGlut2-cre mice. (F) Confocal images representing the expression of tdTomato (red) and mSyp1-eGFP (green) in the LHA. Blue, DAPI. Scale bars, 100 μm. (G) DIO-mSyp1-eGFP in the RVM and DIO-PSD95 tagRFP in the LHA were injected in VGlut2-cre mice. (H) Expression of mSyp1-eGFP (green) and PSD95 (red) in the LHA. Scale bars, 100 μm. (I) Confocal airy scan images of close apposition of mSyp1-eGFP (green) and PSD95-TagRFP (red) at 40x magnification.

Simultaneously, we expressed tdTomato in the Vglut2+ve LHA neurons, which included the LHA_post-RVM_ neurons (Figure 2E). When we imaged the LHA, we found that the GFP-puncta expressed in the synaptic boutons of the glutamatergic RVM neurons surrounded the tdTomato+ve cell bodies of the glutamatergic LHA neurons (Figure 2F). This indicates that the VGlut2+ve RVM neurons synapse with the VGlut2+ve LHA neurons. Further, we tested if the mSyp1-eGFP labeled terminals of the VGlut2+ve RVM neurons appose the post-synaptic components of the dendrites of the VGlut2+ve LHA neurons. The post-synaptic densities in the dendritic compartments of neurons receiving excitatory inputs are rich in PSD95 (Cho et al., 1992). Close apposition between red fluorescent proteins such as tagRFP tagged to PSD95 expressed in the post-synaptic neurons and synaptophysin-tagged GFP in the pre-synaptic neurons have been instrumental in anatomically labeling synaptic connections (Quillet R., 2023; Barik et al., 2018). Thus, in addition to the mSyp1-eGFP expressed in the VGlut2+ve RVM neurons, we expressed PSD95-tagRFP in the VGlut2+ve LHA neurons (Figure 2G). When we imaged the fluorescence in the LHA, we found eGFP-filled synaptic puncta apposing PSD95-tagRFP expressing post-synaptic densities (Figure 2H-I). From this data, we concluded that the excitatory neurons in the RVM form synapses on the LHA excitatory neurons.

### Acute stress activates LHA_post-RVM_ neurons

The glutamatergic neurons in the LHA are stimulated by anxiogenic stimuli such as restraint stress (Owens-French et al., 2022; Wang et al., 2023; Shah et al., 2024) and noxious thermal stimuli (Siemian et al., 2021). Hence, we wondered what stimuli engage the LHA_post-RVM_ neurons. We performed the fiber photometry technique to measure neural activity from the LHA_post-RVM_ neurons in vivo (Figure 3B). In fiber photometry, genetically encoded fluorescent calcium sensors such as GCaMP6s are expressed in targeted neurons, and the calcium dynamics in changing fluorescence in behaving mice are collected through fiber-optic cannulae connected to a CMOS camera (Gunaydin et al., 2014; Kim et al., 2016). In our experiments, we expressed the GCaMP6s (Zhang et al., 2023) in the LHA_post-RVM_ neurons using the transsynaptic genetic labeling strategy described in the previous sections (Figure 3A-C). We found that when mice were placed on the thermal-plate test, innocuous plate temperatures such as 32°C did not elicit a response from the LHA_post-RVM_ neurons. However, when mice were exposed to the 52°C thermal plate temperatures (Espejo and Mir, 1993; Menéndez et al., 2002; Reddy et al., 2023), the activity of the LHA_post-RVM_ neurons gradually rose (Figure 3D). This rise in neural activity can be due to the noxious surface temperature or the stress caused by the mice being in an enclosed, inescapable chamber on a hot surface. Since the rise in activity was gradual, compared to the immediate rise observed in third-order nociceptive neurons in the *Tacr1*-expressing lateral parabrachial nucleus (LPBN) neurons (Barik et al., 2021; Ke et al., 2024), we hypothesized that the LHA_post-RVM_ neurons are preferentially tuned to anxiogenic stimuli. Indeed, when we correlated the neural activity with nocifensive behaviors, we found that the activity of the LHA_post-RVM_ neurons did not correspond to the licks and shakes on the hot plate test (Figure 3E-I). Further, when we applied mildly stressful stimuli to the mice, the LHA_post-RVM_ neurons responded to the bouts of tail hanging and hand immobilization (Figure 3J-Q). Thus, we concluded that the LHA_post-RVM_ neurons are not nociceptive but are tuned to stressful or anxiogenic stimuli.

**Figure 3:**
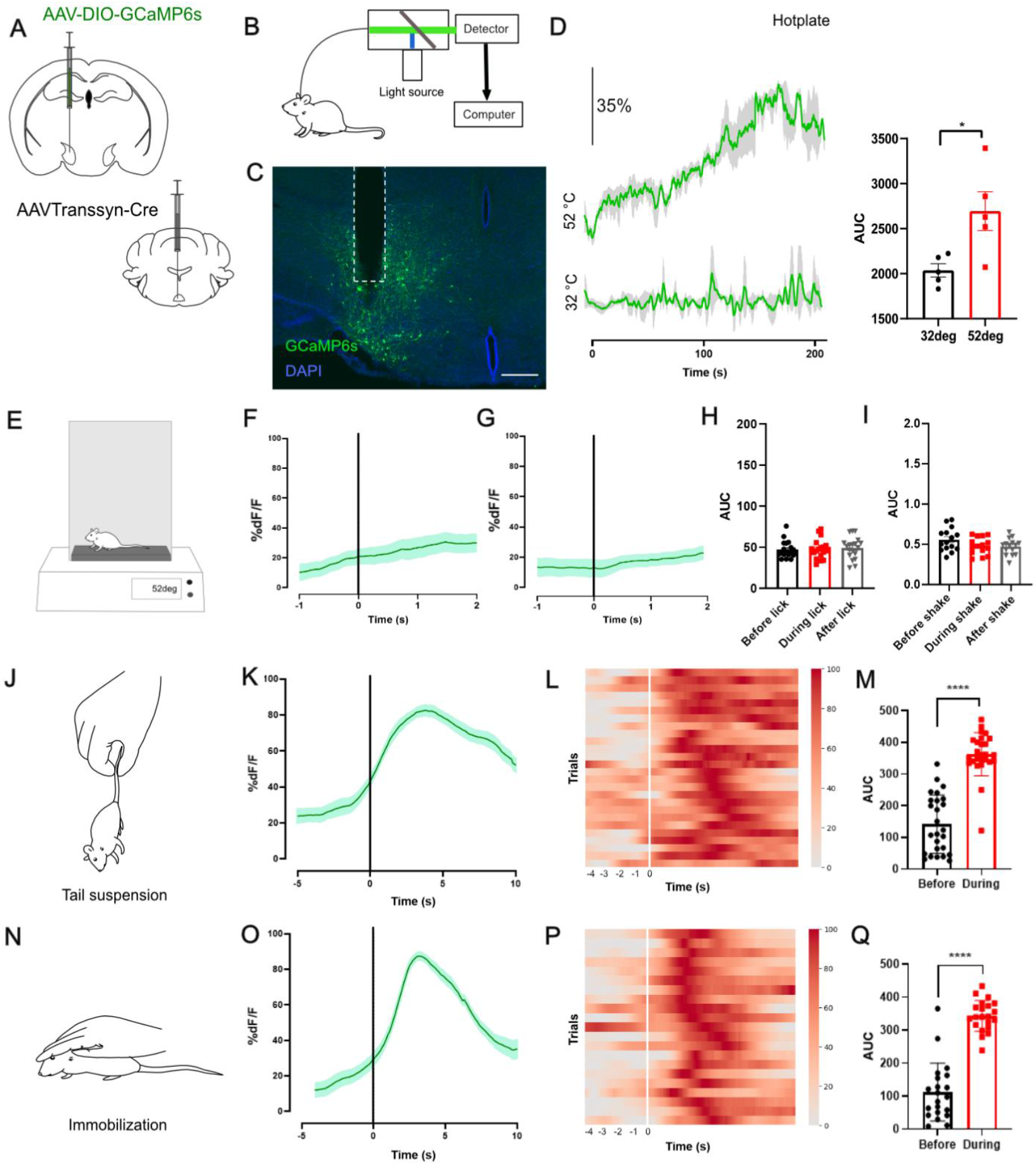
LHA_post-RVM_ neurons respond to mild stressors. (A) AAVTranssyn-Cre in the RVM and DIO-GCaMP6s in the LHA were injected in wild-type mice. (B) General schematics of in vivo fiber photometry recording from mice. (C) Representative image of GCaMP6s expression (green) in the RVM; the white dashed line represents the fiber track. Blue, DAPI. Scale bars, 100 μm. (D) Neural activity traces represent an increase in the overall calcium activity and the area under the curve (AUC) of LHA_post-RVM_ neurons against 52°C and 32°C hotplate (2112.6±118 compared to 2784±190.8, t-test, *p=0.0172, n=5) (E) Mice were subjected to a hotplate at 52°C. (F-G) Mean calcium transients during events (F) licks and (G) shakes (n=5). (H-I) Comparison of AUC: before, during, and after the event. (J) Tail suspension was performed to induce stress in mice. (K) Average calcium transients before and after tail suspension. (L) Heatmap depicting the neural activity of LHA_post-RVM_ neurons for tail suspension trials (n=4; 26 trials). (M) AUC of the neural activity before and during tail suspension (141.65±18.07 compared to 362±13.39, t-test, ****p<0.0001, n=4). (N) Mice were immobilized to induce stress. (O) Average calcium transients for immobilization trials. (P) Heatmap depicting changes in neural activity during immobilization (n=4; 21 trials). (Q) AUC of the neural activity before and during immobilization (112.08±19.11 compared to 343.31±10.18, t-test, ****p<0.0001, n=4).

### Chemogenetic activation and inactivation of LHA_post-RVM_ neurons do not affect nociceptive thresholds

Transient activation of sub-populations of LHA neurons can bi-directionally control nociceptive thresholds (Siemian et al., 2019, 2021; Shah et al., 2024). Hence, we tested if chemogenetic activation and inactivation of the LHA_post-RVM_ neurons would alter the thermal nociceptive thresholds. To attain targeted chemogenetic manipulations of the LHA_post-RVM_ neurons, we expressed the excitatory DREADD (Krashes et al., 2011; Roth, 2016), hM3Dq, and inhibitory DREADD, hM4Di by injecting Cre-dependent AAVs carrying the chemogenetic actuators in the LHA, and AAVTrans-Cre in the RVM. First, we performed the chemogenetic inhibition experiments. Levels of hM4Di expression in the LHA_post-RVM_ neurons were ascertained by visualizing the co-expressed mCherry, which is fused with the hM3Dq at the C-terminus (Figure 4A-B). Intraperitoneal (i.p.) administration of Desclozapine (DCZ) (Nagai et al., 2020) was used to inhibit the hM4Di expressing LHA_post-RVM_ neurons, and saline (vehicle) was used as control. When mice are placed on the hot plate test set at 52°C for 30 seconds, they lick and shake their paw as the surface temperature is in the noxious range and causes thermal pain (Figure 4C). The shakes are spinal reflexes in response to noxious heat, and licks are affective-motivational coping responses under the control of supraspinal circuit mechanisms (Huang et al., 2019; Ma, 2022). We found that inhibition of the LHA_post-RVM_ neurons did not alter the latency or the frequency of lick or shake on the hot plate test (Figure 4E). We then tested the effect of LHA_post-RVM_ inhibition on the tail-flick test, which measures the nociceptive threshold for spinal reflex withdrawal of mouse tail from a source of noxious heat (Irwin et al., 1951) (Figure 4D). We found that chemogenetic inhibition of LHA_post-RVM_ neurons has no bearing on the threshold for the tail-flick response (Figure 4F). Next, we chemogenetically activated the LHA_post-RVM_ neurons by transsynaptically expressing hM3Dq and confirmed the expression by visualizing the hM3Dq-fused mCherry (Figure 4G-H). Neural activation induces the expression of immediate early genes (IEGs) such as Arc and cFos (Morgan et al., 1987; Morgan and Curran, 1989). We found that chemogenetic stimulation of the LHA_post-RVM_ neurons induced cFos expression (Figure 4I). When we tested the effect of activation of the LHA_post-RVM_ neurons on the reflexive and supraspinal behaviors on the hot-plate and tail-flick tests, we found that LHA_post-RVM_ neurons are not sufficient to alter the thresholds or the frequency of occurrence of the nocifensive behaviors (Figure 4J-K). Together, our chemogenetic manipulation experiments indicate that the LHA_post-RVM_ neurons are not involved in determining thermal nociceptive thresholds.

**Figure 4:**
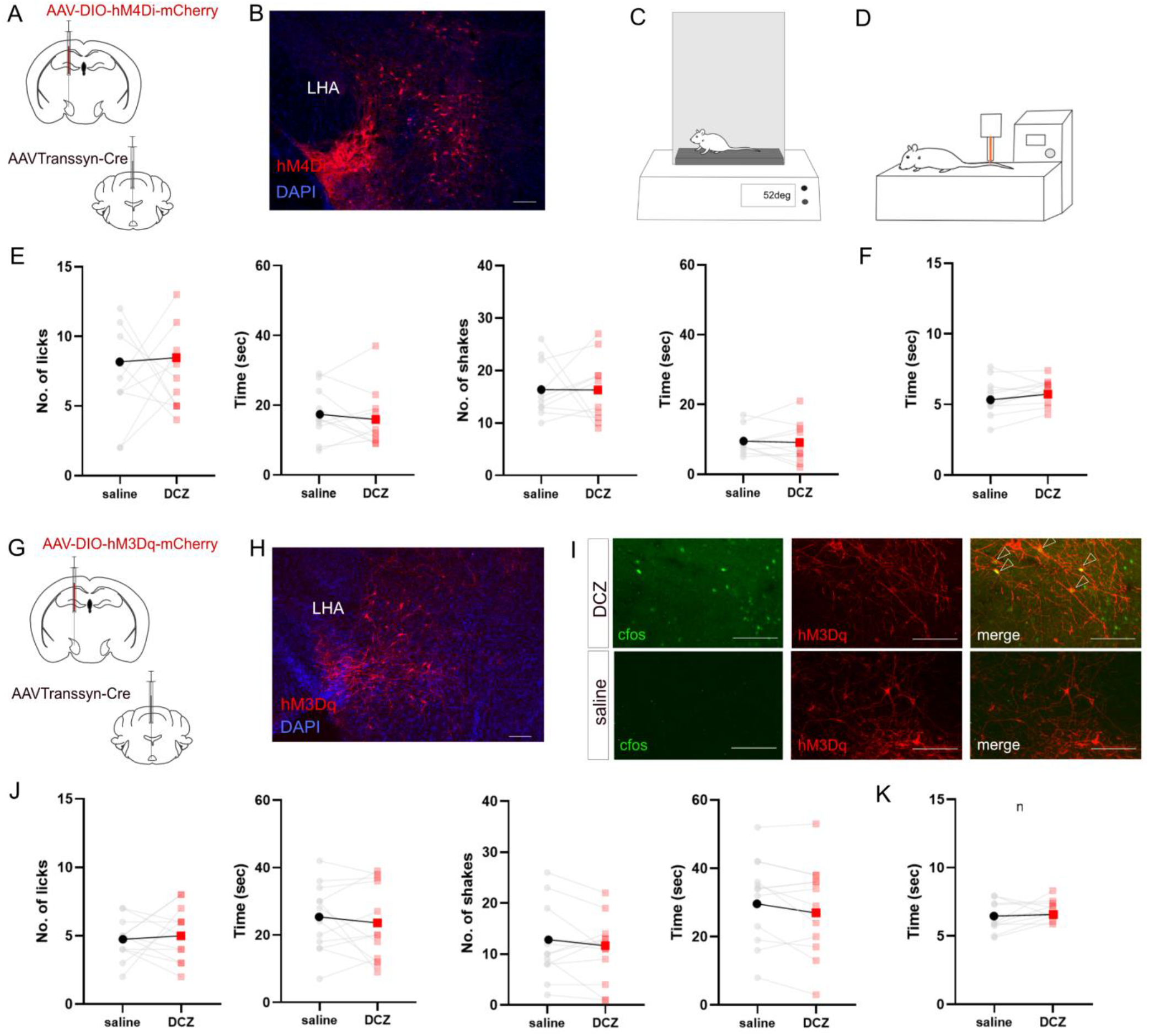
Chemogenetic modulation of LH_post-RVM_ neurons does not affect thermal nociceptive thresholds. (A) AAVTranssyn-Cre in the RVM and DIO-hM4Di in the LHA were injected in wild-type mice. (B) Expression of hM4Di-mCherry (red) in the LHA. (C) Schematics of hotplate assay at 52°C. (D) Schematics of Tail-flick assay. (E) Inhibition of LHA_post-RVM_ neurons does not exhibit any changes in pain behavior when subjected to hotplate and tail-flick: left to right, number of licks, lick latency (seconds), number of shakes, and shake latency (seconds) were recorded. (F) Tail-flick latency (seconds), latency to flick the tail was recorded. (G) AAVTranssyn-Cre in the RVM and DIO-hM3Dq in the LHA were injected in wild-type mice. (H) Expression of hM3Dq-mCherry (red) in the LHA. (I) DCZ induced cFos expression (green) in the LHA. The merged image shows the overlap of cFos expression and hM3Dq (red) expressing LHA_post-RVM_ neurons. (J) Activation of LHApost-RVM neurons does not exhibit any changes in pain behavior when subjected to hotplate and tail-flick: left to right, number of licks, lick latency (seconds), number of shake,s and shake latency (seconds) were recorded. (K) Tail-flick latency (seconds) and latency to flick the tail were recorded. Blue, DAPI. Scale bars, 100 μm.

### RVM-LHA circuitry is sufficient to facilitate aversion and anxiolytic behavior

Aversive, including anxiogenic stimuli, has been shown to engage LHA neurons (Schwartzbaum and Leventhal, 1990; Lecca et al., 2017), in agreement with our findings on the LHA_post-RVM_ neurons. Thus, we tested if the activation of the LHA_post-RVM_ neurons would cause stress and anxiety. A light-dark box is a commonly used behavioral test to determine if laboratory mice are stressed and anxious (Bourin and Hascoët, 2003). In this behavioral paradigm, there are two chambers, one lighted and another dark. When anxious, mice prefer to spend more time in the dark box. We chemogenetically activated the LHA_post-RVM_ neurons by transsynaptically expressing hM3Dq and delivering DCZ i.p. (Figure 5A). Saline-administered mice served as controls. We found that the mice in which LHA_post-RVM_ neurons were chemogenetically activated, spent more time in the dark chamber than the controls indicating that the activation of the LHA_post-RVM_ neurons is anxiogenic (Figure 5B-C). In addition, we tested the effects of DCZ on anxiety levels in mice by transsynaptically expressing GFP in the LHA_post-RVM_ neurons and testing the mice in the light-dark box (Figure 5D). I.p. saline and DCZ resulted in similar time spent in the dark box for the mice expressing GFP in the LHA_post-RVM_ neurons (Figure 5E-F). Further, we asked if RVM_pre-LHA_ drives this anxiolytic behavior in mice. Thus, we injected AAVRetro-Cre in the LHA and Cre-dependent hM3Dq in the RVM (Figure 5G). We found the mice to exhibit similar anxiolytic behavior upon DCZ i.p. administration, as they preferred the dark chamber over the lighted chamber (Figure 5H-C).

**Figure 5:**
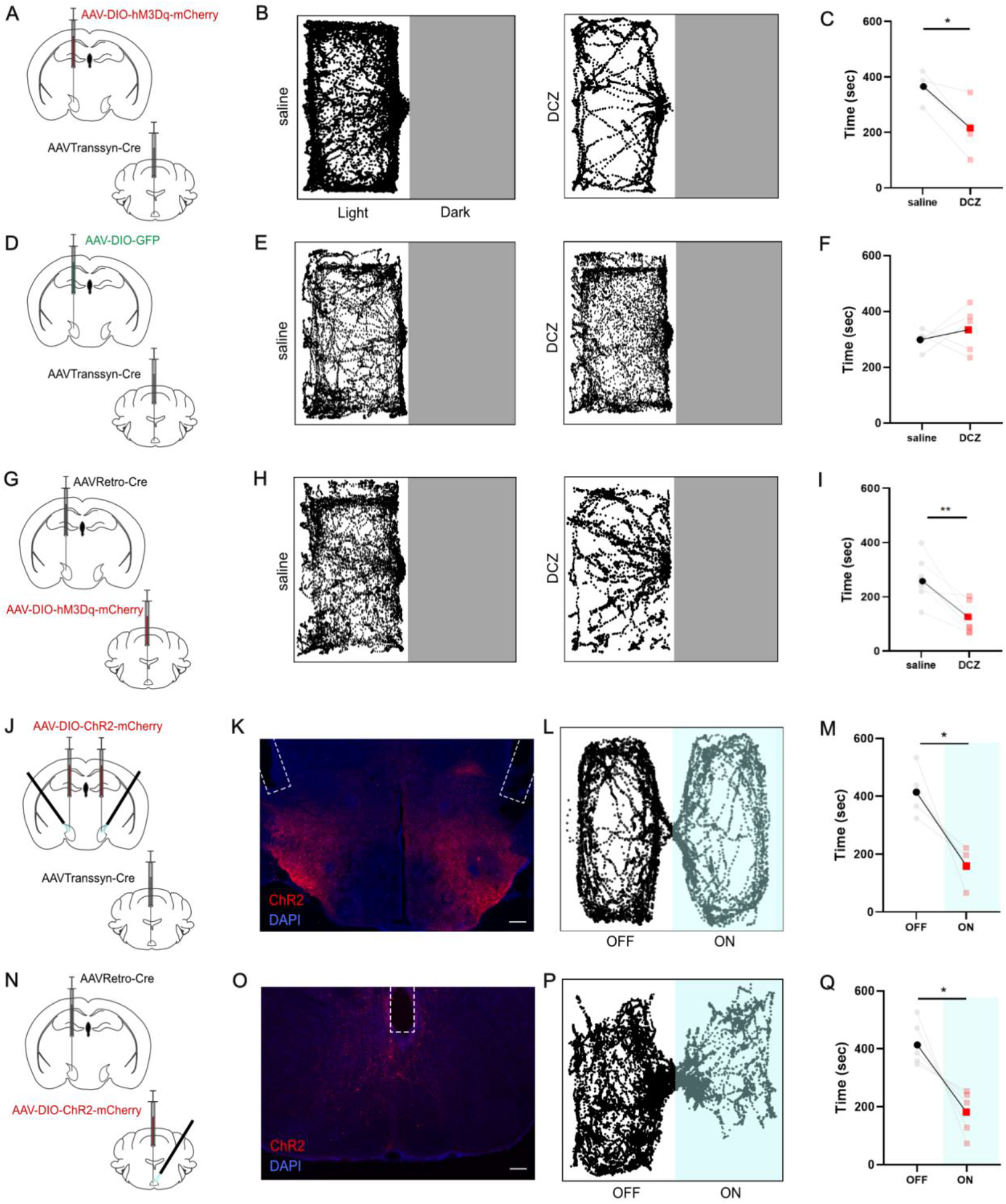
Activation of LH_post-RVM_ neurons causes anxiety-like behaviors. (A) Schematic representation of AAVTranssyn-Cre in the RVM and DIO-hM3Dq in the LHA of wild-type mice. (B) Track plot of mice in the light-dark box representing time spent in the lit chamber after saline and DCZ (i.p.) injection. (C) Upon activation of LH_post-RVM_ neurons, mice spend less time in the light box than saline-treated mice (365.25±27.98 compared to 212.75±50.05, t-test,\**p*=0.026, n=4). (D) Schematic representation of AAVTranssyn-Cre in the RVM and DIO-GFP (control) in the LHA of wild-type mice. (E) Tracking plot of mice in the light-dark box after saline and DCZ (i.p.) injection. (F) Upon DCZ administration in control mice, no changes in time spent in the light box were observed. (G) Schematic representation of AAVRetro-Cre in the LHA and DIO-hM3Dq in the RVM of wild-type mice. (H) Track plot of mice in the light-dark box representing time spent in the lit chamber after saline and DCZ (i.p.) injection. (I) Upon activation of RVM_pre-LHA_ neurons, mice spend less time in the lightbox than saline-treated mice (266.42±41.58 compared to 118.57±23.65, t-test,\*\**p*=0.0034, n=7). (J) AAVTranssyn-Cre in the RVM and DIO-ChR2 in the LHA were injected in wild-type mice. (K) ChR2-mCherry expression in the LHA, where the white dashed line represents the bi-lateral fiber track. Red, ChR2-mCherry; Blue, DAPI. Scale bar, 100 μm. (L) Track plot of RTPA in light-ON and light-OFF chambers. (M) Time spent in the light-ON chamber was significantly less than in the light-OFF chamber (415.5±45.98 compared to 161.25±33.97 t-test, *p=0.048, n=4). (N) DIO-ChR2 in the RVM and AAVRetro-Cre in the LHA were injected in wild-type mice. (O) ChR2-mCherry expression in the RVM, where the white dashed line represents the fiber track. Red, ChR2-mCherry; Blue, DAPI. Scale bar, 100 μm. (P) Track plot of RTPA in light-ON and light-OFF chambers. (Q) Time spent in the light-ON chamber was significantly less than in the light-OFF chamber. (418.0±34.99 compared to 182.0±34.99, t-test, *p=0.028, n=5)

Next, we tested if acute stimulation of the LHA_post-RVM_ neurons is aversive. To that end, we tested if optogenetic activation of the LHA_post-RVM_ neurons would cause aversion in the Real-Time Place Avoidance (RTPA) assay (Lazaridis et al., 2019). The RTPA assay is a two-chambered behavioral test, one of which is paired with optogenetic stimuli. If the optogenetic stimulation of a group of neurons is aversive, it causes avoidance from the paired chamber. We expressed the excitatory optogenetic actuator, Channelrhodopsin-2 (ChR2) (Zhang et al., 2006; Deisseroth and Hegemann, 2017), with the transsynaptic strategy in the LHA_post-RVM_ neurons (Figure 5J-K). We found that the mice expressing ChR2 in the LHA_post-RVM_ neurons avoided the chamber in which blue light was delivered in LHA via optic cannulae upon entrance (Figure 5L-M). Thus, transiently activating LHA_post-RVM_ neurons is sufficient to cause aversion and avoidance. Next, we wondered if similar optogenetic activation of the RVM neurons pre-synaptic to the LHA_post-RVM_ (RVM_pre-LHA_) neurons would avoid the optogenetically paired chamber in the RTPA assay. In order to express ChR2 in the RVM_pre-LHA_ neurons, we injected AAVRetro-Cre in the LHA and intersected with the Cre-dependent DIO-ChR2-mCherry in the RVM (Figure 5N). By visualizing the fluorescence, we confirmed the successful labeling of the RVM_pre-LHA_ neurons with ChR2-mCherry (Figure 5O). When we activated the RVM_pre-LHA_ neurons by shining blue light on entering the paired chamber in the RTPA assay, we found that the RVM_pre-LHA_ neurons were sufficient to drive avoidance behavior (Figure 5P-Q). Together, we conclude that either the pre-synaptic RVM_pre-LHA_ or the post-synaptic LHA_post-RVM_ neurons mediate acute aversion and avoidance behaviors.

### RVM_pre-LHA_ neurons are nociceptive

We found that the LHA_post-RVM_ neurons do not respond to noxious somatosensory stimuli yet are actively involved in the aversive aspect of pain (Figure 3D). Hence, we sought to understand whether pre-synaptic RVM_pre-LHA_ neurons are tuned to noxious somatosensory stimuli. Many RVM neurons are pro-nociceptive and thus respond to high-threshold mechanical and thermal stimuli (Fields and Basbaum, 1978; Basbaum and Fields, 1984; Nguyen et al., 2023). We performed fiber photometry recordings from the RVM_pre-LHA_ neurons to determine the stimuli they respond to. We expressed the GCaMP6s in the RVM_pre-LHA_ neurons by stereotaxically delivering AAVRetro-Cre bilaterally in the LHA and DIO-GCaMP6s in the RVM (Figure 6A-B). It was found that the RVM_pre-LHA_ neurons, similar to the LHA_post-RVM_ neurons, respond to stressful stimuli. When we hung the mice by their tail for 2 seconds, we found that the activity in the RVM_pre-LHA_ neurons rose while the mice were hanging, and gradually went down when the mice were returned to their homecage (Figure 6C). This suggests that the RVM_pre-LHA_ neurons respond to anxiogenic stimuli. Next, we tested if the RVM_pre-LHA_ neurons respond to noxious heat on the spinal reflex tail-flick test (Figure 6D). In the tail-flick test, the activity of the RVM_pre-LHA_ neurons increased post-withdrawal, indicating that the neural activity builds up as mice are exposed to noxious heat stimuli (Figure 6E-H). We found that mechanical stimuli (tail pinch) engage RVM_pre-LHA_ neurons too; thus, these neurons’ activity is not solely tuned to the noxious thermal stimuli (Figure 6I-L). On the gradient hot plate test (32 to 52°C, 5 mins), the RVM_pre-LHA_ neurons were active around 40°C when the temperature entered the noxious range (Figure 6M-O). This indicates that the RVM_pre-LHA_ neurons are tuned to noxious thermal stimuli and can be instrumental in driving the nocifensive responses to thermal pain.

**Figure 6:**
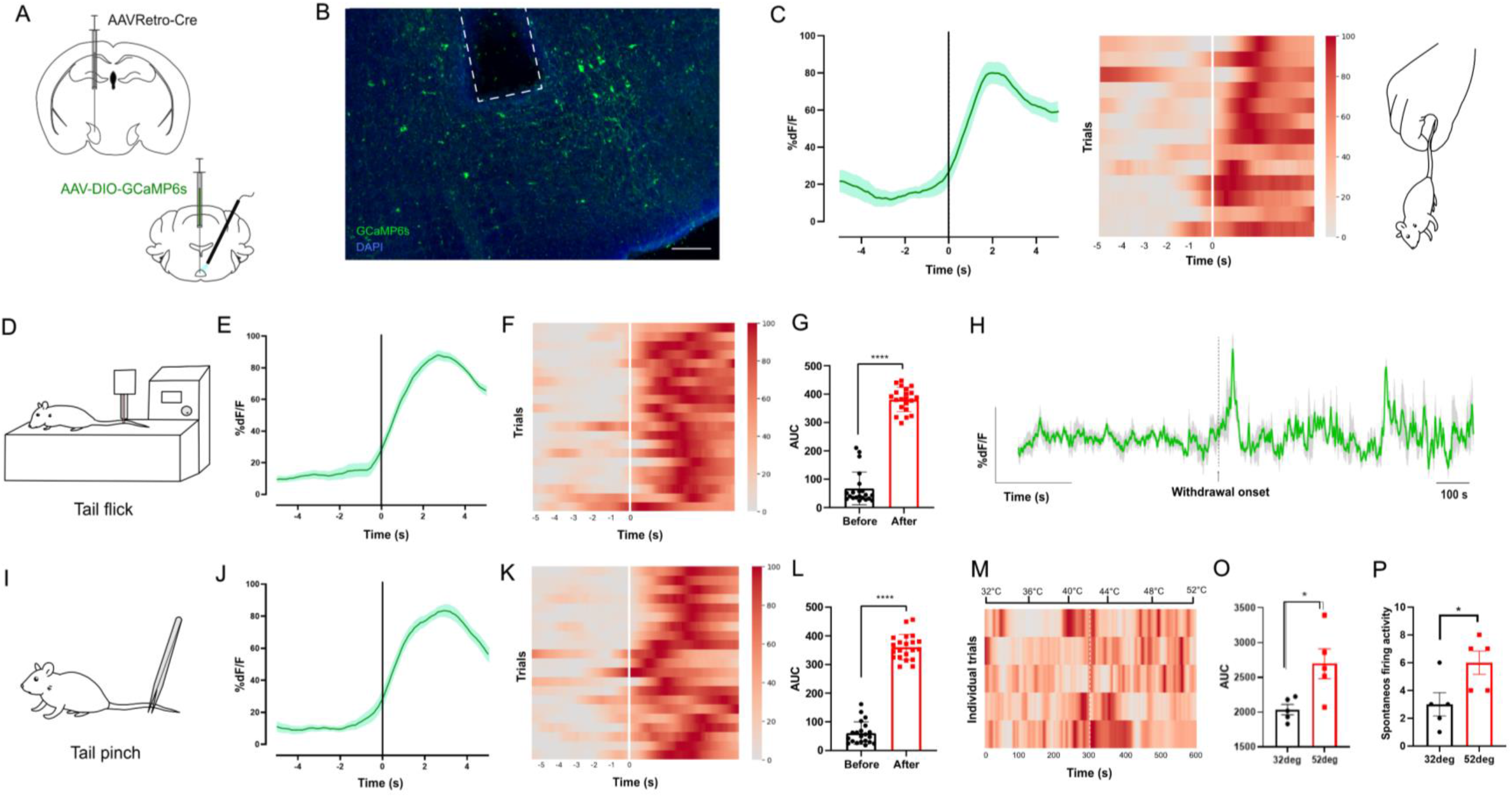
RVM_pre-LHA_ neurons are nociceptive. (A) In wild-type mice, AAVRetro-Cre in the LHA and DIO-GCaMP6s in the RVM were injected. Fiber photometry recording was done from the RVM. (B) A representative image of the GCaMP6s expression (green) in the RVM, with a white dashed line representing the fiber track. Blue, DAPI. Scale bars, 100 μm. (C) Cumulative plot and heat map for calcium transients when mice were suspended from the tail (n=4, 13 trials). (D) Schematic of tail-flick assay. (E) Mean calcium transients for tail flick trials (n=5). (F) Heatmap depicting neural activity in RVM_pre-LHA_ neurons before and after tail-flick response (n=5; 21 trials). (G) The area under the curve (AUC) of the neural activity before and during the behavioral trial (67.12±13.24 was compared with 371.21±11.73, t-test, ****p<0.0001, n=5). (H) Sample cumulative trace of overall neural activity when subjected to tail-flick. (I) Schematic of tail pinch assay. (J) Mean calcium transients of LHA_post-RVM_ neurons during tail pinch (n=5; 21 trials). (K) Heatmap depicting neural activity before and after a tail pinch. (L) Area under the curve (AUC) of the neural activity before and during the behavioral trial (65.12±12.34 was compared with 384.21±9.40, t-test, ****p<0.0001, n=5). (M) Heatmap representing neural activity during gradient hotplate, 32°C to 52°C (n=5). (N) The AUC was found to be significantly increased at 52°C vs 32°C (2037±74.13 compared to 2695±216.5, t-test, *p=0.0207, n=5). (O) The spontaneous firing activity of LH_post-RVM_ neurons was found to be significantly increased at 52°C vs 32°C (3.00±0.837 compared to 6±0.837, t-test, *p=0.0350, n=5).

Thus, RVM_pre-LHA_ neurons are engaged by anxiogenic and noxious thermal somatosensory stimuli.

### Brain-wide mapping of the LHA_post-RVM_ neuronal projections

The spinal projections of the brainstem structure RVM have been studied in terms of their role in pain modulation. However, the axonal arborizations of the RVM neurons to the forebrain and midbrain structures remain poorly explored. To this end, we found that the RVM neurons project to the midbrain LHA and play an essential role in the affective-motivational components of pain, such as stress and anxiety. Here, we sought to map the brain-wide projections of the LHA_post-RVM_ neurons. To that end, we injected AAVTranssyn-Cre in the RVM and DIO-GFP in the LHA of wild-type mice to express GFP in the LHA_post-RVM_ neurons (Figure 7A). After three weeks of viral infection, we observed robust GFP expression at the injection site in the LHA (Figure 7B).

**Figure 7:**
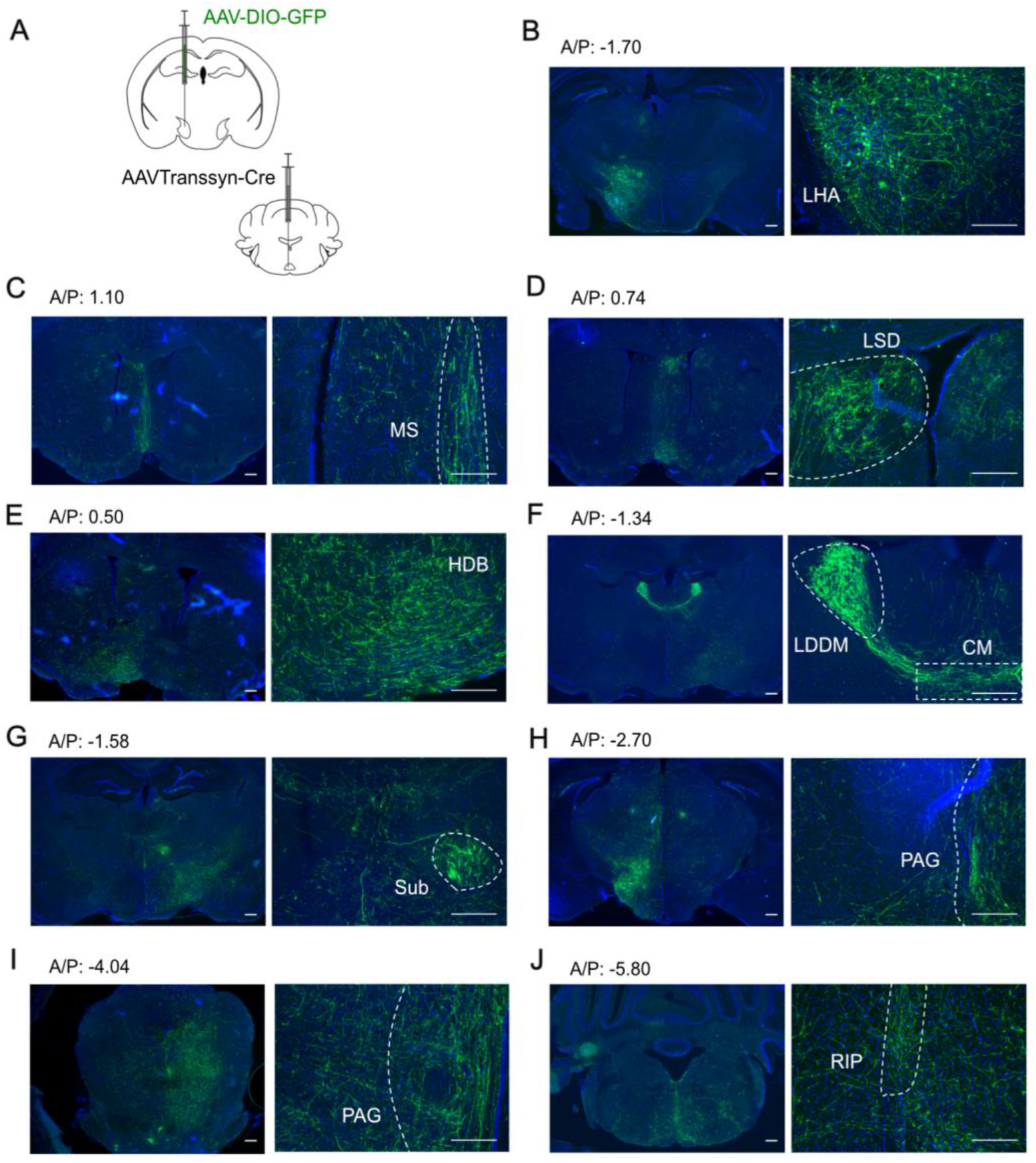
Axonal projections of LHA_post-RVM_ neurons. (A) Schematics for mapping downstream targets by injecting AAVTranssyn-Cre in the RVM and DIO-GFP in the LHA of wild-type mice. (B) The site of injection at the LHA shows GFP-positive neurons. (C-J) Axon projections of LHA_post-RVM_ neurons representing downstream targets. LHA, Lateral Hypothalamic area; RVM, Rostral Ventromedial Medulla; MS, Medial Septal nu; LSD, Lateral Septal nu; HDB, nu Horizlimb Diagonal Band; LDDM, Laterodorsal thalamic nu (dorsomedial); CM, Central Medial thalamic nu; Sub, Submedius thalamic nu; PAG, Periaqueductal gray; RIP, Raphe Interpositus nu. Green, eGFP; Blue, DAPI. Scale bars, 100 μm.

Dense GFP expressing axon terminals of the LHA_post-RVM_ neurons were observed in the forebrain regions such as medial septum (MS), dorsal lateral septum (LSD), and central medial thalamic nuclei (CM), known to be involved in the affective-motivational components of pain (Figure 7C-F). Furthermore, we observed the LHA_post-RVM_ axonal terminals at the submedial thalamic nuclei (Sub) and the periaqueductal gray (PAG), essential for driving nociceptive and defensive behaviors (Figure 7G-I). PAG is well-known for initiating the freezing response when subjected to fear conditioning (Koutsikou et al., 2014). In contrast, Sub is a region comprising nociceptive neurons involved in the ventrolateral orbital cortex (VLO)-mediated descending antinociceptive pathway (Tang et al., 2009). Other LHA_post-RVM_ targets, such as the horizontal diagonal band (HDB), laterodorsal thalamic nuclei (LDDM), and raphe interpositus nuclei (RIP) of the brain-stem, can provide novel insights into our understanding of pain and stress-induced anxiety (Figure 7J).

## Discussion

How noxious stimuli are transformed into stress is poorly understood. RVM, a brainstem nuclei rich in the site of action of endogenous opioids through their projections to the spinal cord, are known to respond to and bi-directionally control pain thresholds. However, it was not known whether RVM neurons play a role in the emotional effects of pain through their brain projections. Here, we show that the RVM neurons project to the LHA, a critical node for stress-modulation of pain, to drive anxiety-like behaviors. Interestingly, the LHA downstream neurons did not affect nociceptive thresholds. Thus, we found a novel role for the nociceptive neurons in the RVM, which, through their LHA projections, can mediate the anxiogenic effects of pain.

The LS-LHA-RVM descending pathway has been shown to mediate stress-induced analgesia (Shah et al., 2024). We have now tested the role of an ascending neural pathway between RVM and LHA in thermal nociception and anxiety. Our anatomical data indicates that the LS and RVM neurons synapse onto distinct LHA populations (Figure 1), indicating that they have independent functions. Indeed, LHA_post-RVM_ neurons, unlike LHA_post-LS,_ do not modulate nociceptive thresholds; however, they can drive anxiogenic behaviors (Figure 5). Thus, the excitatory neurons in the LHA, depending on the inputs, can differentially mediate pain and stress-induced anxiety. This can be explained by the fact that the LHA_post-RVM_ neurons, unlike LHA_post-LS,_ do not project to the RVM but rather to nuclei such as the medial septum and central-medial thalamic nuclei, which play instrumental roles in the affective-motivational behaviors.

The RVM consists of the midline nucleus raphe magnus (NRM) and the adjacent nucleus reticularis gigantocellularis of the reticular formation. The NRM consists of serotonergic neurons simultaneously releasing enkephalins and serotonin (Potrebic et al., 1994; Wei et al., 2010). These neurons have been exclusively studied in pain and itch modulation, owing to their projections to the superficial dorsal horn (Liu et al., 2014). It will be interesting to test the role of the ascending projections of the brain projections of the NRM serotonergic neurons exclusively and whether the neurons that project to the spinal cord also project to the brain or not. The RVM neurons have been functionally divided into ON, OFF, and Neutral cells (Fields et al., 1991). The ON cells respond to noxious stimuli and are inhibited by opioid administration (Fields et al., 1983; Barbaro et al., 1986). In contrast, the OFF neurons are inhibited by noxious stimuli, and opioids have an excitatory effect on these cells. The RVM-neutral cells, as the name suggests, are indifferent toward noxious stimuli and opioids. Several serotonergic neurons were found to be OFF cells (Potrebic et al., 1994). It was found that the RVM serotonergic neurons do not participate in opioid-mediated analgesia (Gao et al., 1998). Thus, the RVM serotonergic neurons might have independent functions of the neurons involved in opioidergic pain modulation.

Together, we have delineated a novel neural pathway between RVM and LHA that may play an essential role in the affective-motivational components of pain. Future studies will reveal if the RVM_pre-LHA_ and the LHA_post-RVM_ neurons are involved in the stress and anxiety seen in humans and animals with chronic pain.

